# Repertoire analysis of antibody CDR-H3 loops suggests affinity maturation does not typically result in rigidification

**DOI:** 10.1101/230417

**Authors:** Jeliazko R. Jeliazkov, Adnan Sljoka, Daisuke Kuroda, Nobuyuki Tsuchimura, Naoki Katoh, Kouhei Tsumoto, Jeffrey J. Gray

**Author notes:** These authors contributed equally to this work (Jeliazko R. Jeliazkov, Adnan Sljoka). **Correspondence:** Adnan Sljoka or Jeffrey J. Gray.

## Abstract

Antibodies can rapidly evolve in specific response to antigens. Affinity maturation drives this evolution through cycles of mutation and selection leading to enhanced antibody specificity and affinity. Elucidating the biophysical mechanisms that underlie affinity maturation is fundamental to understanding B-cell immunity. An emergent hypothesis is that affinity maturation reduces the conformational flexibility of the antibody’s antigen-binding paratope to minimize entropic losses incurred upon binding. In recent years, computational and experimental approaches have tested this hypothesis on a small number of antibodies, often observing a decrease in the flexibility of the Complementarity Determining Region (CDR) loops that typically comprise the paratope and in particular the CDR-H3 loop, which contributes a plurality of antigen contacts. However, there were a few exceptions, and previous studies were limited to a small handful of cases. Here, we determined the structural flexibility of the CDR-H3 loop for thousands of recently-determined homology models of the human peripheral blood cell antibody repertoire using rigidity theory. We found no clear delineation in the flexibility of naïve and antigen-experienced antibodies. To account for possible sources of error, we additionally analyzed hundreds of human and mouse antibodies in the Protein Data Bank through both rigidity theory and B-factor analysis. By both metrics, we observed only a slight decrease in the CDR-H3 loop flexibility when comparing affinity-matured antibodies to naïve antibodies, and the decrease was not as drastic as previously reported. Further analysis, incorporating molecular dynamics (MD) simulations, revealed a spectrum of changes in flexibility. Our results suggest that rigidification may be just one of many biophysical mechanisms for increasing affinity.

## 1 Introduction

Antibodies are proteins produced by the B cells of jawed vertebrates that play a central role in the adaptive immune system. They recognize a variety of pathogens and induce further immune response to protect the organism from external perturbation. Molecules that are bound by antibodies are referred to as antigen and are recognized by the antibody variable domain (Fv), which is comprised of a variable heavy (V_H_) and light (V_L_) domain. To overcome the challenge of recognizing a vast array of targets — the number of antigens being far greater than the number of antibody germline genes — antibodies rely on combinatoric and genetic mechanisms that increase sequence diversity (1–3). Starting from a limited array of germline genes, a naïve antibody is generated by productive pairing of a randomly recombined V_H_, assembled from V-, D-, and J-genes on the heavy locus, and randomly recombined V_L_, assembled from V- and J-genes on the kappa and lambda loci (1). Next, in a process known as affinity maturation, iterations of somatic hypermutation are followed by selection to evolve the antibody in specific response to a particular antigen. This evolution results in the gradual accumulation of mutations across the entire antibody, with higher mutation rates in the six complementarity determining regions (CDRs) than in the framework regions (FRs) (4, 5). The CDRs are hypervariable loops comprising a binding interface on the Fv domain beta-sandwich framework, with three loops contributed by each chain; the light chain CDRs are denoted as L1, L2, and L3 and the heavy chain CDRs are H1, H2, and H3. The five non-H3 CDRs can be readily classified into a discrete amount of canonical structures (6–10) because they possess limited diversity in both sequence and structure. The CDR-H3 on the other hand is the focal point of V(D)J recombination, resulting in exceptional diversity of both structure and sequence. While all CDRs contribute to antigen binding, the diverse CDR-H3 is often the most important CDR for antigen recognition (11–15). Thus, to understand the role of B cells in adaptive immunity and how they evolve antibodies capable of binding specific antigens, we must first understand the effects of affinity maturation on the CDRs, and in particular on the CDR-H3.

Over the last 20 years, the structural effects of affinity maturation have been studied with an assortment of experimental and computational methods. X-ray crystallography has been used to compare antigen-inexperienced (naïve) and antigen-experienced (mature) antibodies with both antigen present and absent. Analysis of the catalytic antibodies 48G7, AZ-28, 28B4, and 7G12 showed a 1.2 Å average increase in Cα RMSD of the CDR-H3 upon antigen binding in the naïve over that of the mature antibody, whereas motion in the other CDRs varied (16–20). Beyond structural studies, surface plasmon resonance (SPR) has been used to assess the energetics and association/dissociation rate constants of antibody-antigen binding. Manivel ***et al.*** studied a panel of 14 primary (naïve) and 11 secondary (mature) response anti-peptide antibodies, observing that affinity maturation resulted in increases in the association rate and corresponding changes in the entropy of binding (21). Schmidt ***et al.*** saw the opposite when studying a broadly neutralizing influenza virus antibody, observing that affinity maturation resulted primarily in a decrease in the dissociation rate, with little effect on the association rate (22). Isothermal calorimetry (ITC) has also been used to determine antigen-binding energetics including the enthalpic and entropic contributions. For nine anti-fluorescein antibodies, including 4-4-20 and eight anti-MPTS antibodies, ITC results revealed diverse effects of affinity maturation: 14 of 17 mature antibodies bound antigen in an enthalpically favorable and entropically unfavorable manner, yet 3 of 17 showed the opposite, with entropically favorable and enthalpically unfavorable binding energetics (23, 24). Three-pulse photon echo peak shift (3PEPS) spectroscopy has been used to quantify dynamics of chromophore-bound antibodies on short timescales of femto- to nanoseconds. 3PEPS spectroscopy results from a panel of 18 antibodies showed that mature antibodies can possess a range of motions from small rearrangements such as side-chain motions to large rearrangements such as loop motions (23–25). In a specific comparison of naïve vs. mature, for the 4-4-20 antibody, the mature antibody was found to have smaller motions, i.e. to be more rigid, than naïve (23–28). Antibody dynamics have also been studied by hydrogen-deuterium exchange mass spectroscopy (HDX-MS), which in contrast to 3PEPS probes timescales of seconds to hours. Comparison of three naïve and mature anti-HIV antibodies showed changes in CDR-L2/H2, but not in CDR-H3 dynamics (29). Finally, MD simulations have been used to study antibody dynamics on intermediate timescales of nano- to microseconds. MD simulations showed rigidification and reduction of CDR-H3 loop motion upon maturation for seven studied naïve/mature antibodies, with two exceptions, depending on the specific study (22, 28, 30-34). In an orthogonal protein design approach to examine the CDR-H3 loop flexibility, Babor ***et al*** and Willis ***et al.*** found that naive antibody structures are more optimal for their sequences, when considering multiple CDR-H3 loop conformations (35, 36). In sum, past studies focusing on the effects of affinity maturation on CDRs have found evidence suggesting that mature antibodies have more structural rigidity and less conformational diversity than their naïve counterparts (16, 18, 19, 23–27).

With recent growth in the number of antibody structures deposited in the Protein Data Bank (PDB) and development of homology models from high-throughput sequencing of paired VH-VL genes in B cells, we now have the datasets necessary to test the rigidity hypothesis on a large scale. Prior studies, usually focused on a few antibodies at time, generally support the hypothesis that affinity maturation rigidifies the CDR-H3 loop. Thus, we hypothesize that this effect should be observable in a repertoire-scale study of thousands of antibodies. We first analyzed thousands of recently determined RosettaAntibody homology models of the most common antibody sequences found in the human peripheral blood cell repertoire (37). We estimated the structural flexibility of the CDR-H3 loop by applying the Floppy Inclusions and Rigid Substructure Topography (FIRST) and the Pebble Game (PG) algorithms to determine backbone degrees of freedom (DOFs). Surprisingly, we found no difference in the CDR-H3 loop flexibility of the naïve and mature antibody repertoires. We considered alternative explanations for our results, which were incongruent with past studies, by expanding our analysis to a large set of antibody crystal structures, including several previously characterized antibodies, and extending our methods to include other measures of flexibility such as B-factors and MD simulations. By all analysis methods, we found mixed results: some antibodies’ CDR-H3 loops were more flexible after affinity maturation whereas others’ became less flexible. In summary, we find that while affinity maturation can modulate antibody binding activity by reducing CDR-H3 structural flexibility, it does not necessarily do so.

## 2 Materials and Methods

### 2.1 Immunomic Repertoire Modeling

Briefly, RosettaAntibody is an antibody modeling approach that assembles homologous structural regions into a rough model and then refines the model through gradient-based energy minimization, side-chain repacking, rigid-body docking, and ***de novo*** loop modeling of the CDR-H3. The approach is fully detailed in (38) and (39). In a typical simulation, ~1,000 models are generated and the ten lowest-energy models are retained. The immunomic repertoire we analyzed is from DeKosky and Lungu, ***et al.*** (37). In that study, models were generated for each of the 500 most frequently occurring naïve and mature antibody sequences in two donors (a total ~20,000 models representing the ~2,000 most frequent antibodies).

### 2.2 Structural Rigidity Determination

The flexibility or rigidity of the CDR-H3 loop backbone was determined by using several extensions of the Pebble Game Algorithm (PG) (40–43) and method FIRST (44); we refer to here as FIRST-PG. For a given protein structure, FIRST generates a molecular constraint network consisting of nodes (atoms) and edges (interactions representing covalent bonds, hydrogen bonds, hydrophobics etc.). Each potential hydrogen bond is assigned an energy in kcal/mol which is dependent on donor-hydrogen acceptor geometry. FIRST is run with a selected hydrogen-bonding energy cutoff, where all bonds weaker than this cutoff are ignored in the network. On the resulting network, the PG algorithm is then used to identify rigid clusters, flexible regions, and overall available conformational degrees of freedom (DOFs). For a given antibody structure, DOFs for the protein backbone of the CDR-H3 loop were calculated at every hydrogen-bonding energy cutoff value between 0 to −7 kcal/mol in increment steps of 0.01 kcal/mol. This calculation was repeated for every member of that antibody ensemble (i.e. ten lowest energy models of the ensemble) and finally, at each energy cutoff, the DOF count was averaged over the entire ensemble. For a given energy cutoff and a given member of the ensemble, the DOF count for the CDR-H3 loop (residues 95-102) was obtained by calculating the maximum number of pebbles that belong to the backbone atoms (Cα, C, N) of the CDR-H3 loop (40).

### 2.3 Degree of Freedom Scaling

To compare flexibility across CDR-H3 loops of different lengths, the DOF metric computed above is scaled by a theoretical maximum DOF. We define 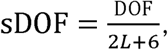 where, *2L* (the loop length in residues) represents the backbone degrees of freedom (torsion angles: ϕ,ψ), and *6* represents the trivial but ever-present rigid-body DOFs (rotations/translations in 3D).

### 2.4 Area Under Curve Calculation

The area under the curve (AUC) is approximated by simple numerical integral (akin to trapezoidal integration), where the first term defines a rectangle and the second term defines a triangle:

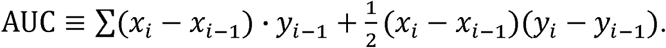

### 2.5 Crystallographic Dataset

On June 27^th^, 2017, a summary file was generated from the Structural Antibody Database (SAbDab) (45), using the “non-redundant search” option to search for antibodies with maximum 99% sequence identity, paired heavy and light chains, and a resolution cutoff of 3.0 Å. The summary file, containing 1021 antibodies, was used as input to a SAbDab download script which yielded corresponding sequences, Chothia-numbered PDBs, and IMGT data (on occasion this had to be updated to match the reported germline in the IMGT 3Dstructure-DB) (46). The structures were further pruned: structures were omitted if there were unresolved CDR-H3 residues, as this would preclude flexibility calculations, or if the antibody was neither human nor mouse, as this would prevent alignment to germline. Prior to analysis, structures were truncated to the Fv region (removing all residues but light chain residues numbered 1-108 and heavy chain residues numbered 1-112, in Chothia numbering) and duplicate and non-antibody (for example, bound antigen) chains were removed. A total of 922 antibody crystal structures were analyzed. The following CDR definitions were used throughout this paper, in conjunction with the Chothia numbering scheme: L1 spans light chain residue numbers 24–34, L2 spans 50-56, L3 spans 89-97, H1 spans heavy chain residue numbers 26-35, H2 spans 50-56, and H3 spans 95-102.

### 2.6 Alignment to Germline

The germline of each antibody was determined by IMGT lookup (46) Then, BLASTP (version 2.2.29+) with the BL0SUM50 scoring matrix was used to align the antibody variable region heavy and light sequences to corresponding germline sequences (IGHV, IGKV, and IGLV loci only, downloaded from IMGT). The number of mismatches according to BLAST were considered as the number of amino acid mutations from germline. Supplementary Table 1 details the PDB ID, CDR-H3 length, number of heavy chain mutations, number of light chain mutations, heavy germline gene, and light germline gene data for each structure in the dataset.

### 2.7 B-Factor Z-Score Calculation

Temperature factors (B-factors) were extracted for all Cα atoms in the variable region of the antibody heavy chain (V_H_, Chothia numbering 1-112). The arithmetic mean and sample standard deviation values were calculated for the B-factors. For each Cα atom in the CDR-H3 region, residue numbers spanning 95-102 under the Chothia numbering convention (11), the z-score was calculated as 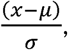 where *x* is the B-factor of the current Cα atom and *μ* and *σ* are the mean and standard deviation of B-factors for all Cα atoms in the V_H_ respectively.

### 2.8 Rosetta Relaxation And Ensemble Generation

Antibody structural ensembles with 10 members were generated using either the Rosetta FastRelax (47, 48) or Rosetta KIC protocol (49). The Rosetta FastRelax protocol consists of five cycles of side-chain repacking and gradient-based energy minimization in the REF2015 version of the Rosetta energy function (50). Thus, FastRelax ensembles explore the local energy minimum of the crystal structure. The KIC ensembles are more diverse and representative of RosettaAntibody homology models: each ensemble member was generated by running the CDR-H3 refinement step of the RosettaAntibody protocol, consisting of V_H_-V_L_ docking, CDR-H3 loop remodeling, and all-CDR loop minimization (38, 39). Sample command lines are given in the Supplementary Material. The structural ensembles produced by both FastRelax and KIC were used for rigidity analysis.

### 2.9 Molecular Dynamics Simulations

The Fv regions were retrieved from the original PDB files. The MD simulations were performed using the NAMD 2.12 package (51) with the CHARMM36m force field and the CMAP backbone energy correction (52). The truncated Fv structures were solvated with TIP3P water in a rectangular box such that the minimum distance to the edge of the box was 12 Å under periodic boundary conditions. Na or Cl ions were added to neutralize the protein charge, then further ions were added corresponding to a salt solution of concentration 0.14 M. The time step was set to 2 fs throughout the simulations. A cutoff distance of 10 Å for Coulomb and van der Waals interactions was used. Long-range electrostatics were evaluated through the Particle Mesh Ewald method (53).

The initial structures were energy-minimized by the conjugate gradient method (10,000 steps), and heated from 50K to 300K during 100 ps, and the simulations were continued by 1 ns with NVT ensemble, where protein atoms were held fixed whereas non-protein atoms freely moved. Further simulations were performed with NPT ensemble at 300K for 200 ns without any restraints other than the SHAKE algorithm to constrain bonds involving hydrogen atoms. The last 180 ns of each trajectory was used for the subsequent clustering analyses. Similar to a previous work (54), a total of 2000 evenly spaced frames from each trajectory were clustered based on root-mean-square deviation (RMSD) of the Cα and Cβ atoms using the K-means clustering algorithm implemented in the KCLUST module in the MMTSB tool set (55). The cluster radius was adjusted to maintain 20 clusters in each trajectory. The structure closest to the center of each cluster was chosen as a representative structure of each cluster. The 10 representative structures were chosen from the top 10 largest clusters and these representative structures were energy-minimized by the conjugate gradient method (10,000 steps) in a rectangular water box. The minimized antibody Fv structures were used as the inputs for the rigidity analysis.

Root-mean-square quantities of the MD trajectories were calculated based on the last 180 ns trajectories. After superposing Cα atoms of the FR of the heavy chain (FR_H_) of each snapshot onto Cα atoms of FR_H_ of the reference structures (i.e. crystal structures), Cα-RMSD of CDR-H3 was calculated as the time average. Similarly, after superposing Cα atoms of entire Fv domains of each snapshot onto those of the reference structures, the root-mean-square fluctuation (RMSF) of a residue *i* was defined as the time average:

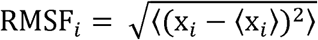

where x_i_ is the distance between the Cα atom of the snapshots at a given time and the Cα atom of the *i*th residue of the reference structures (56).

## 3 Results

### 3.1 Immunomic Repertoire Reveals No Difference in Flexibility between Naïve and Mature CDR-H3 Loops

We initially asked whether CDR-H3 loop rigidification, having been observed in many past studies, was present in a large set of antibodies derived from human peripheral blood cells. Previously, DeKosky and Lungu *et al.* used Rosetta Antibody to model the structures of ~2,000 common antibodies found in the peripheral blood cells of two human donors (37). Paired V_H_–V_L_ sequences were derived from either CD3^-^CD19^+^CD20^+^CD27^-^ naïve B cells or CD3^-^CD19^+^CD20^+^CD27^+^ antigen experienced B cells (mature) isolated from peripheral mononuclear cells. RosettaAntibody structural models were created by identifying homologous templates for the CDRs, V_H_-V_L_ orientation, and FRs; assembling the templates into one model; *de novo* modeling the CDR-H3 loop; rigid-body docking the VH-VL interface; side-chain packing; and minimizing in the Rosetta energy function (38). Since *de novo* modeling of long loops is challenging, DeKosky and Lungu *et al.* limited their antibody set to the more tractable subset of antibodies with CDR-H3 loop lengths under 16 residues. They compared their models for seven human germline antibodies with solved crystal structures and found models had under 1.4 Å backbone RMSD for the FR and under 2.4 Å backbone RMSD for the CDR-H3 loop.

We used the FIRST-PG method (40, 44) to estimate flexibility from the RosettaAntibody homology models, determining the number of backbone DOFs for the CDR-H3 loop as each hydrogen bond is broken in order from weakest to strongest. FIRST models the antibody as a molecular graph where nodes represent atoms and edges represent atomic interactions. An extension of the PG algorithm uses this molecular graph to compute the DOFs of the CDR-H3 loop. To mitigate the effects of homology modeling inaccuracies on the FIRST-PG analysis, we used an ensemble of ten lowest-energy RosettaAntibody models. FIRST-PG analysis on structural ensembles has been shown to predict hydrogen-deuterium exchange and protein flexibility (51). To account for varying CDR-H3 loop lengths, we scaled the calculated DOFs by a theoretical maximum value (Methods). Figure 1A shows a curve of the scaled DOFs averaged over all naïve or mature antibodies as a function of the hydrogen-bonding energy cutoff used in the FIRST-PG analysis. At a cutoff of 0 kcal/mol, all hydrogen bonds are intact and the average CDR-H3 loop scaled DOFs are about 20% of the theoretical maximum. Moving from right to left on the plot increases the minimum energy cutoff for including interactions in the FIRST graph; effectively hydrogen bonds of increasing strength are “broken” and the available DOFs rise from 20% to over 90% of the maximum theoretical flexibility while the loop becomes unstructured (unfolded) in FIRST.

**Figure 1.**
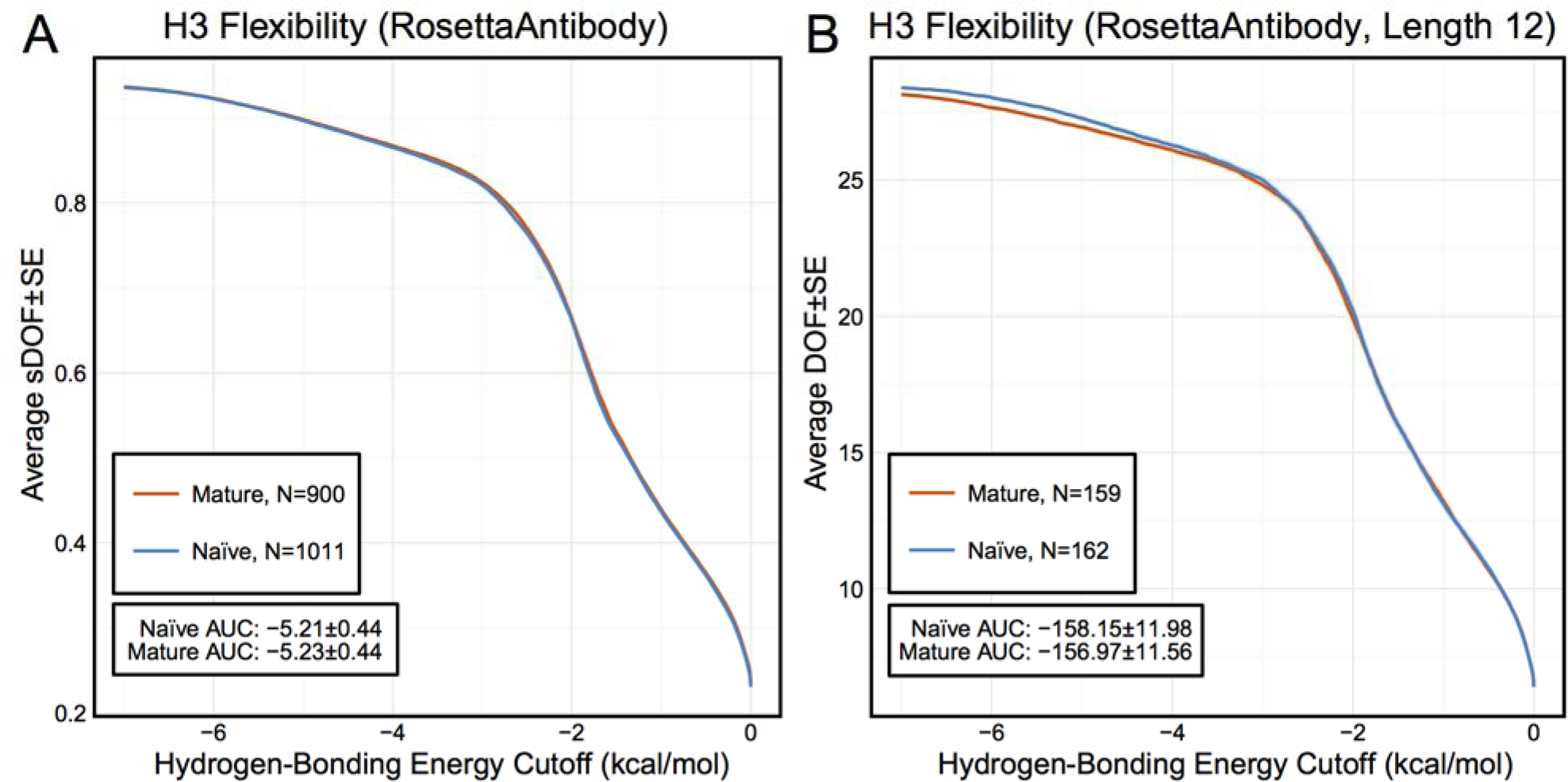
FIRST-PG analysis of the immunomic antibody set created by Rosetta Antibody modeling, with naïve antibody data shown in blue and mature antibody data shown in orange and standard error of the mean shown in a lighter shade of the respective color. FIRST-PG analysis calculates the DOFs of CDR-H3 loop as a function of hydrogen-bonding energy cutoff. (A) When comparing DOFs scaled to a theoretical maximum as a function of hydrogen-bonding energy cutoff for the entire set, the values are similar for both naïve (AUC ± SD = −5.2 ± 0.44) and mature (AUC ± SD = −5.2 ± 0.44) antibodies. (B) Comparison of DOFs for a single length without scaling yields similar results, compare the naïve AUC ± SD at −158.15 ± 11.98 and mature at −156.97 ± 11.56.

In comparing the curves for naïve and mature antibodies (Figure 1A), there is no difference in the average, scaled DOFs. To quantify this comparison, we computed the average AUC plus-or-minus one standard deviation for both antibody sets. The average AUC values are identical between naïve (-5.21 ± 0.44) and mature antibody repertoires (-5.23 ± 0.44). This lack of difference persists (AUC −158.15 ± 11.98 [naïve] vs. −156.97 ± 11.56 [mature]) when accounting for CDR-H3 loop length (Figure 1B), and so the observed similarity of DOFs in naïve and mature antibodies is not due to averaging over loops of different lengths. Thus, on the immunomic repertoire scale, we do not observe the difference in flexibility between naïve and mature antibodies predicted by the paratope rigidification hypothesis.

Before amending the rigidification hypothesis in light of these results, we considered several alternative explanations for our observations. First, we addressed whether the use of homology models for flexibility analysis introduced inaccuracies by analyzing a large set of antibody crystal structures and Rosetta-generated models from that set with varying quality, ranging from models with sub-angstrom backbone RSMD to models that may be several angstroms off (and more representative of an average homology model). Next, we addressed whether backbone DOFs, as calculated by FIRST-PG, were a good measure of flexibility, by assessing flexibility through two alternative measures: B-factors and MD simulations. Additionally, we addressed whether averaging flexibilities and comparing across many germlines affected results, by detailed flexibility analysis of previously studied naïve-mature antibody pairs and RosettaAntibody-modeled pairs.

### 3.2 Only Small Flexibility Differences Are Observed Between Naïve and Mature Antibodies in the Crystallographic Set

#### 3.2.1 Preparation of an Antibody Crystal Structure Dataset

Of course, the strongest critique of the immunomic antibody set is that these models are only approximating the actual antibody structure. Thus, we applied FIRST-PG analysis to a large set of antibody crystal structures. We curated the set of all non-redundant mouse and human antibody crystal structures from SAbDab (45). To be consistent with the models produced by RosettaAntibody, we truncated the structure of each antibody to only the Fv domain, excluding other antibody regions or antigen. Then, we used IMGT/3Dstructure-DB (57) to identify the variable domain genes and determined the number of somatic mutations by aligning the sequence derived from the crystal structure to the IMGT-determined gene. We defined mature antibodies as those possessing at least one somatic mutation in either V gene. Our full dataset has 922 antibodies of which 23 are naïve. CDR-H3 loop lengths and germline assignments are summarized in Supplementary Table 1. Summary statistics are plotted in Supplementary Figures 1–3.

#### 3.2.2 FIRST-PG Analysis of Crystal Structures

From the crystal structures, we created two sets of structural ensembles and assessed flexibility by FIRST-PG. Flexibility analysis has previously been shown to be more accurate on ensembles in comparison to analysis using single (snapshot) conformers (41, 58). Ensembles of ten representative structures were generated from the initial crystal structure by using either using Rosetta FastRelax (47) or the refinement step of RosettaAntibody (38, 39), which we term KIC ensembles after the loop modeling algorithm used in refinement (49). Rosetta FastRelax samples structures around the crystallographic, local energy-minimum, with typically < 1 Å backbone RMSD, whereas the refinement step of RosettaAntibody samples a more diverse set of low-energy CDR-H3 loop conformations and Vh-VL orientations. Thus, FastRelax ensembles are representative of the crystal structures, whereas KIC ensembles are representative of RosettAntibody homology models. By comparative FIRST-PG analysis of the two sets, we can assess the effects of modeling inaccuracies on flexibility analysis.

The scaled DOFs as calculated by FIRST-PG for FastRelax ensembles of antibody crystal structures are shown in Figure 2A. There are only minor differences between the naïve and mature flexibility curves and the AUC is similar for both sets (-4.70 ± 0.46 [naïve] vs. −4.70 ± 0.48 [mature]). Again, we considered the possibility that different distributions of loop lengths in the two sets obscures the affinity maturation contributions to flexibility. Therefore, we analyzed loops of length 10 (Figure 2B), the single most common length in our set. When loops of a single length were compared, there was a separation between the naïve and mature sets, with the naïve antibody set average DOFs being consistently greater than the mature set. The AUC values differ, but are within a standard deviation (-128.2 ± 9.0 [naïve] vs. −121.9 ± 10.1 [mature]). We repeated FIRST-PG analysis for KIC ensembles of antibody crystal structures and observed similar results (Supplementary Figure 4): for scaled DOFs, the AUC was −5.91 ± 0.20 (naïve) vs. −5.81 ± 0.26 (mature) and, for loops of length 10 only, the AUC was −154.10 ± 4.80 (naïve) vs. −150.44 ± 7.73 (mature). Thus, there does not appear to be a large, consistent CDR-H3 loop flexibility difference across all antibodies, but rather there is a small difference for antibodies with similar-length CDR-H3 loops.

**Figure 2.**
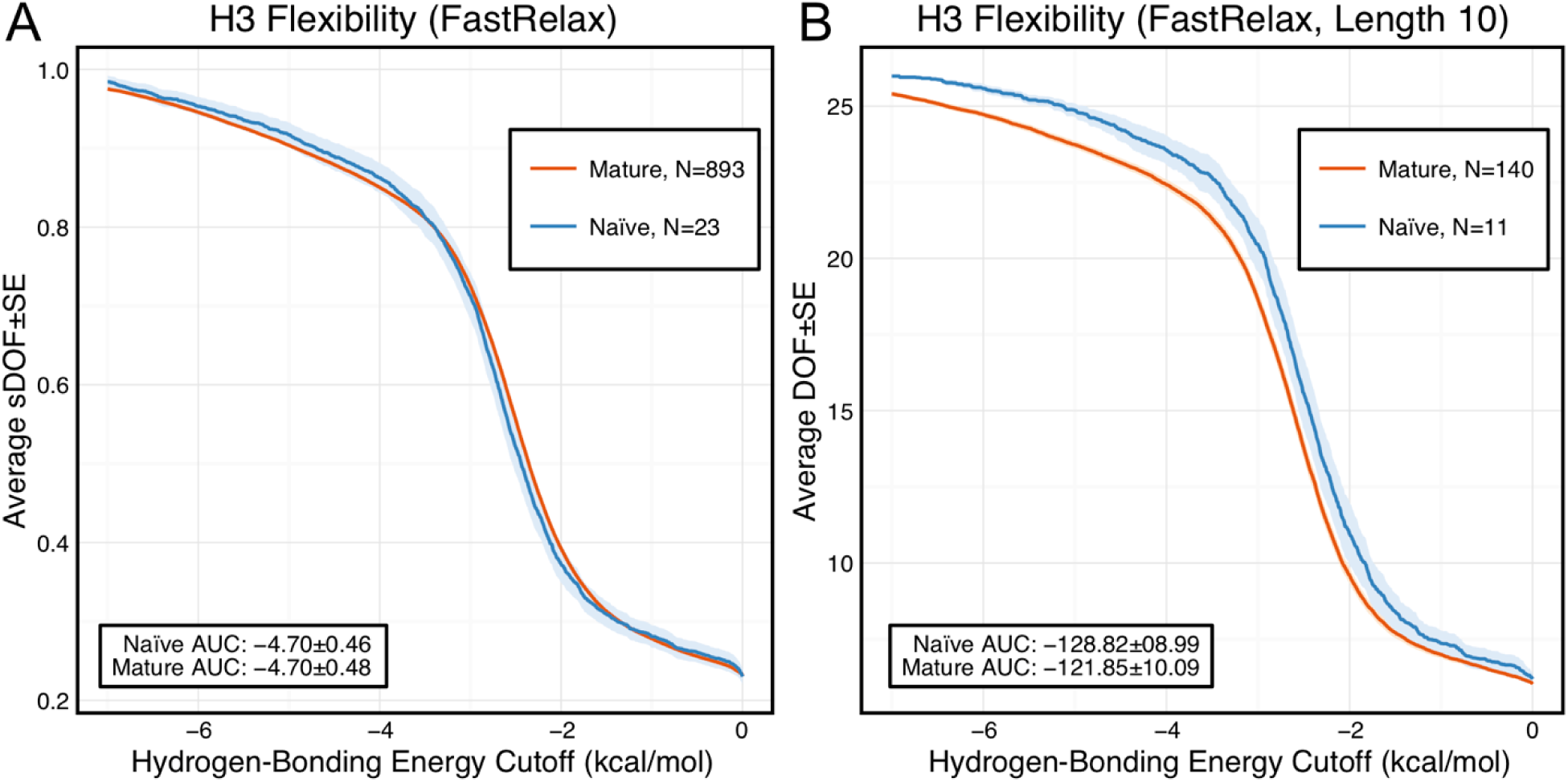
FIRST-PG analysis of the crystallographic antibody set, with naïve antibody data shown in blue and mature antibody data shown in orange and standard error of the mean shown in a lighter shade of the respective color. (A) When comparing DOFs scaled to a theoretical maximum as a function of hydrogen-bonding energy cutoff for the entire set, the values are similar for both naïve (AUC = −4.7 ± 0.46) and mature (AUC = −4.7 ± 0.48) antibodies. (B) Comparison of DOFs for a single length without scaling reveals naïve antibodies to possess a slightly higher DOF value than mature antibodies at the same hydrogen-bonding energy cutoff. AUCs however are within a standard deviation, compare naïve at −128.82 ± 8.99 and mature at −121.85 ± 10.09.

#### 3.2.3 B-Factor Analysis of Crystal Structures

However, we have not accounted for the possibility that backbone DOFs as calculated by FIRST-PG may not capture the effects of affinity maturation on CDR-H3 loop flexibility. Thus, we assessed loop flexibility as determined by atomic temperature factors or B-factors. In protein crystal structures, B-factors measure the heterogeneity of atoms in the crystal lattice. Thus, rigid regions have lower B-factors as they are more homogenous throughout the crystal whereas flexible regions have higher B-factors as they are less homogenous throughout the crystal. B-factors are also affected by crystal resolution, so we cannot compare raw values across structures of varying resolution. Instead, we computed a normalized B-factor z-score, which has zero mean and unit standard deviation for each antibody chain. Finally, to account for different CDR-H3 loop lengths, we averaged the B-factor z-scores for the CDR-H3 loop residues.

Figure 3 shows the distributions of B-factor z-scores averaged over the CDR-H3 loop residues of naïve and mature antibodies. Both distributions span a similar range and overlap significantly, with the naïve curve peak shifted toward higher values than the mature. The majority of the naïve CDR-H3 loop B-factor z-score averages were positive (65%), whereas the majority of the mature CDR-H3 loop B-factor z-score averages were negative (64%). A two-sample Kolmogorov-Smirnov (KS) test confirms the distributions to be distinct, with a maximum vertical deviation, *D,* of 0.36 and a p-value of 0.006.

**Figure 3.**
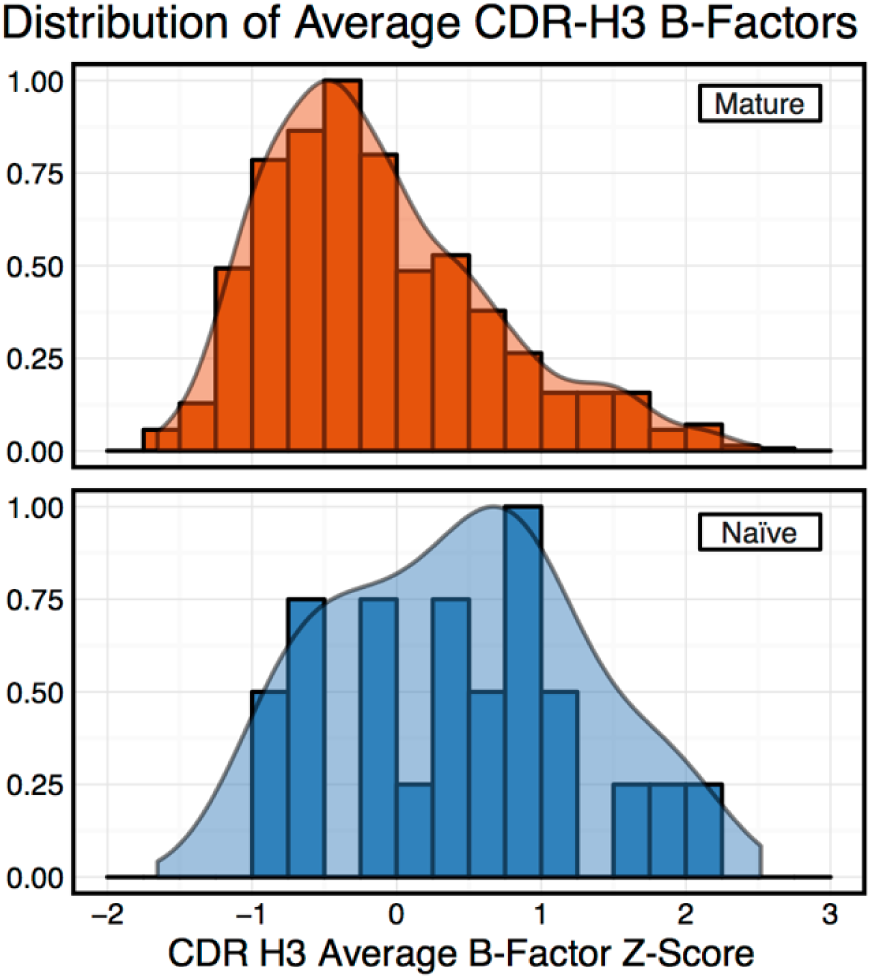
Distributions of CDR-H3 loop average B-factors for the crystallographic set of antibodies. Bars show binned counts in intervals of 0.25, with the maximum bar height scaled to 1, whereas smoothed densities are normalized to integrate to 1. Distributions split by number of somatic mutations appear distinct, despite significant overlap (mature shown in dark orange, naïve shown in dark blue). A two-sample KS test confirms different underlying distributions with a p-value of 0.006 and maximum vertical deviation, D, of 0.36.

However, we were concerned that the mixing of bound and unbound crystal structures would influence results, as we previously observed bound structures to have lower average B-factors (59). Furthermore, in the PDB-derived dataset, naïve antibodies were mostly to be crystallized in the unbound state (19 of 23), whereas mature antibodies were mostly to be co-crystallized with their cognate antigen (544 of 899). In conjunction, these two observations suggested that the high number of antigen-bound mature antibody crystal structures was the primary driver of the difference between naïve and mature B-factor z-scores. Thus, we compared the B-factor averages of unbound structures only and found that while the distributions appear to be distinct (Figure 4), they fail a two-sample KS test (D = 0.27, p = 0.15). As we conjectured, the primary difference was found to be between the bound and unbound distributions (Figure 5), with a two-sample KS test confirming the difference between the distributions (D = 0.31, p < 2. 16E-16). Additionally, we considered other possible origins of difference between the naïve and mature distributions that are not related to affinity maturation, including comparison across species, crystal structure resolutions, CDR-H3 loop lengths, and if the CDR-H3 loop was at a crystal contact or not. We found none of these to have as clear of an effect on the distribution of B-factor averages as whether or not antigen was bound (Supplementary Figures 5 and 6). In summary, the distributions of B-factor z-score averages (Figures 3–5) suggest that both the naïve and mature antibody sets possess CDR-H3 loops of varying flexibility and that neither set is significantly more flexible or rigid than the other.

**Figure 4.**
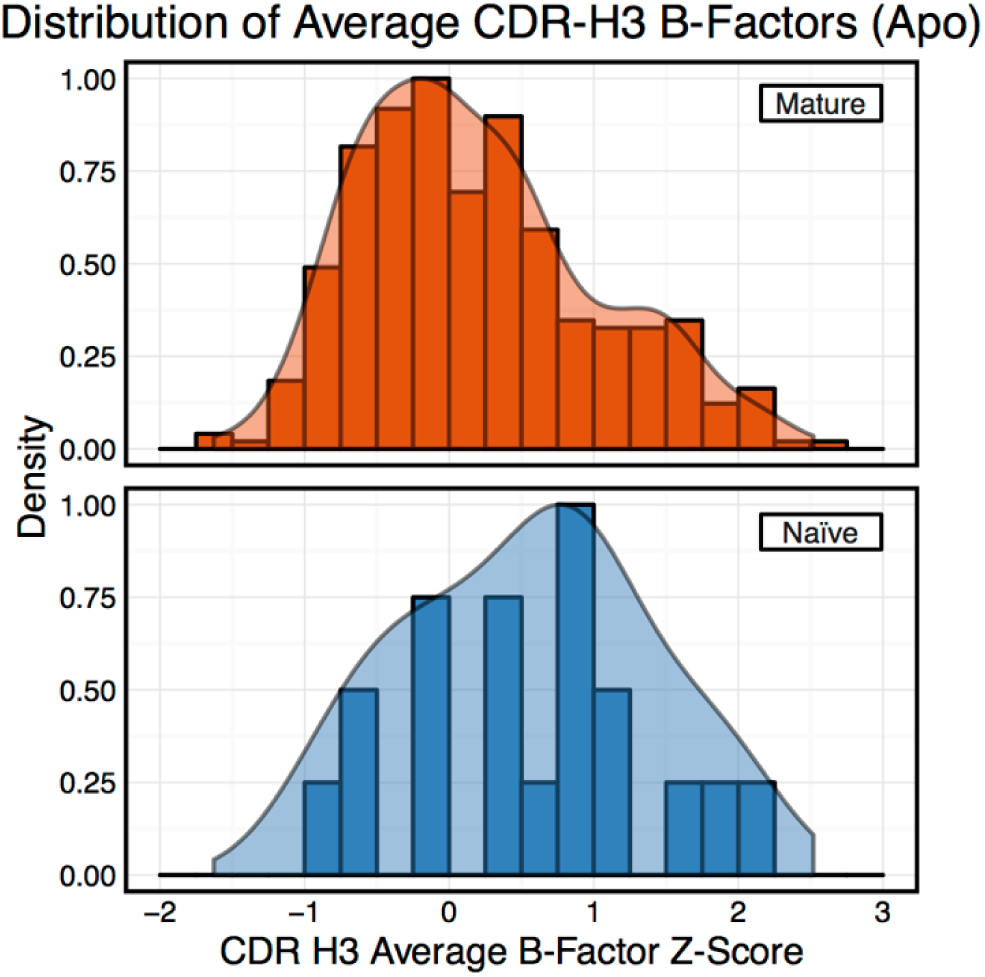
Distributions of CDR-H3 loop average B-factors for the crystallographic set of unbound antibodies. Bars show binned counts in intervals of 0.25, with the maximum bar height scaled to 1, whereas smoothed densities are normalized to integrate to 1. Distributions split by number of somatic mutations appear distinct, despite significant overlap (mature shown in dark orange, naïve shown in dark blue). However, a two-sample KS test indicates identical underlying distributions with a p-value of 0.15 and maximum vertical deviation, D, of 0.27.

**Figure 5.**
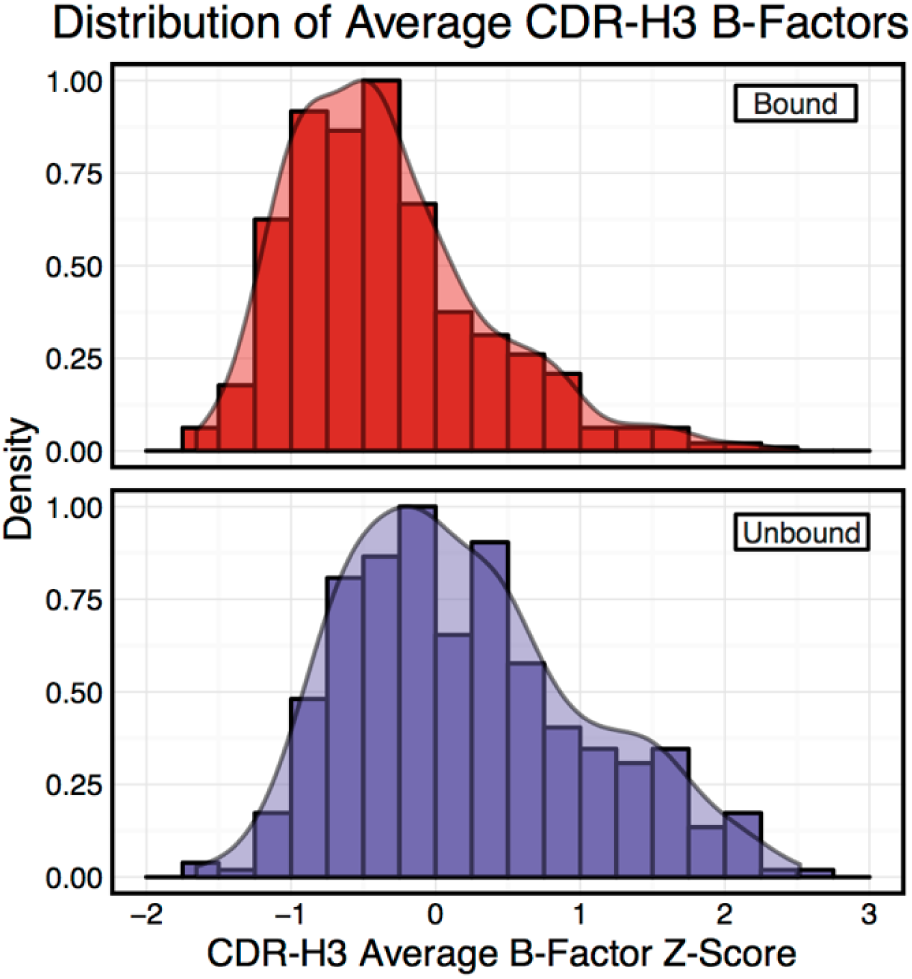
Distributions of CDR-H3 loop average B-factors for the crystallographic set of antibodies. Bars show binned counts in intervals of 0.25, with the maximum bar height scaled to 1, whereas smoothed densities are normalized to integrate to 1. Distributions split by whether or not antigen is present in the crystal structure appear distinct (bound shown in red, unbound shown in purple). A two-sample KS test confirms different underlying distributions with a p-value of 2.2E-16 and maximum vertical deviation, D, of 0.31.

### 3.3 Comparison of Mature to Naïve-Reverted Models Reveals Varying Rigidification Across Matched Pairs

Based on the B-factor results from the 922 analyzed crystal structures, we postulated that rigidification was not a repertoire-wide phenomenon (i.e. all mature antibodies are not more rigid than all naïve antibodies), but it could still be plausible that matched paris of naïve and mature antibodies would reveal rigidification.

To investigate this hypothesis, we selected ten mature antibodies from our SAbDab set with CDR-H3 loops of length 10, a length for which loop modeling performs well (49, 60). To control for species, half of the selected antibodies were human and half were mouse. We reverted the mature antibody sequences to naïve using the germline sequences from the aligned V-genes. We then used RosettaAntibody to generate homology models for the naïve-reverted sequences. We analyzed the ensembles of the ten lowest-energy homology models using FIRST-PG. To ensure fair comparison, we also used FIRST-PG to analyze homology model ensembles of the mature sequences. To provide an estimate for the accuracy of RosettaAntibody homology models, we computed RMSDs for the mature models using the known crystal structures and found all had sub-2-Å CDR-H3 loop backbone RMSD, calculated after alignment of the heavy chain FR, with 4 of 10 antibodies having sub-Å RMSD (Supplementary Figures 7–11).

Of the ten naïve/mature antibody pairs we analyzed, six showed a decrease in flexibility and four showed an increase in flexibility upon affinity maturation (Figure 6). These ten antibodies demonstrate the breadth of possible affinity maturation effects, from an expected flexibility decrease in antibody 2AGJ, with AUC decreasing by 9.34%, to the unexpected flexibility increase in antibody 1RZ7, with AUC increasing by 10.65%.

**Figure 6.**
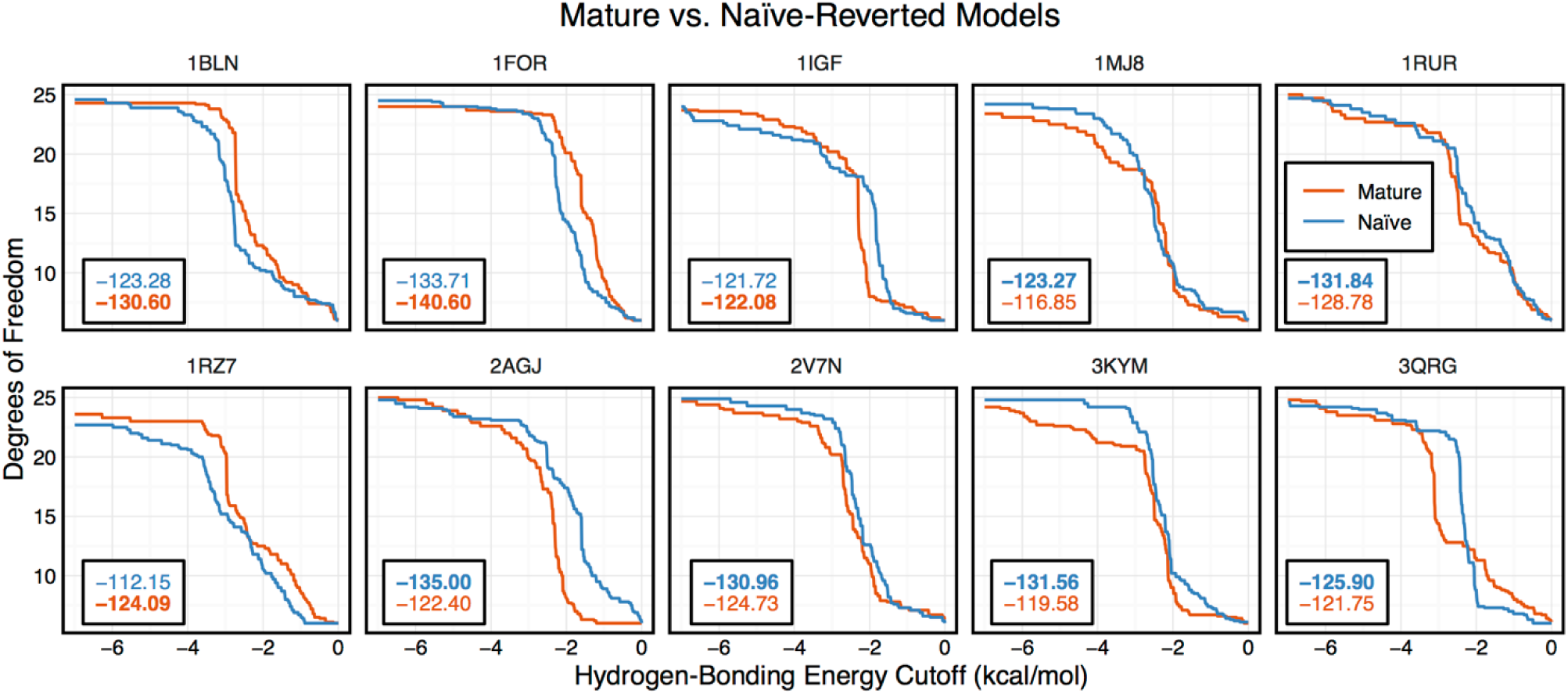
FIRST-PG analysis of ten RosettaAntibody-modeled mature/naïve-reverted antibody pairs (CDR-H3 loop length of 10). Naïve values are colored blue, while mature values are color orange. AUCs are reported in the bottom left of each sub-figure, with bold indicating the greater value. Four out of the ten cases have mature antibodies with AUC greater than their naïve counterparts.

### 3.4 Analysis of 48G7 Antibody

Having analyzed 1911 models, 922 crystal structures, and 10 paired-reverted models, we had yet to observe a consistent difference in CDR-H3 loop flexibility between naïve and mature antibodies, as previously reported in literature. Thus, we turned to three previously studied antibodies with known crystal structures and measured CDR-H3 loop flexibility. These are (1) the esterolytic antibody 48G7 (16, 32, 33, 35), (2) the anti-fluorescein antibody 4-4-20 (23, 26–28, 31, 33), and (3) a broadly neutralizing influenza virus antibody (22). For all three antibodies, the effects of affinity maturation on CDR-H3 loop flexibility have been previously studied by both experiment and simulation, allowing comparison with our results. For brevity, we presently discuss the 48G7 antibody here, and full results for all antibodies are available in the Supplementary Material.

The 48G7 antibody was first studied through crystallography, with structures capturing the bound (holo) and unbound (apo) states of both the naïve and mature antibody (16). Comparison between the naïve and mature CDR loop motions from the free to the bound state revealed minor changes, with the mature CDR-H3 loop being slightly more rigid and moving an Angstrom less than the naïve upon antigen binding (Supplementary Figures 12 and 13). For each of the four crystal structures, we extracted B-factors and computed B-factor z-scores for the CDR-H3 loop, measuring the distance from the B-factor mean in standard deviations. B-factor z-scores for the CDR-H3 loop of apo-48G7 are shown in Figure 7A. The mature antibody has lower B-factors than the naïve antibody throughout the entire CDR-H3 loop. This observation also holds for the holo-48G7 antibody structures as well (Supplementary Figure 14). Supplementary Table 2 summarizes B-factors averaged over the whole CDR-H3 loop. These B-factor results agree with the prior crystallographic observations.

**Figure 7.**
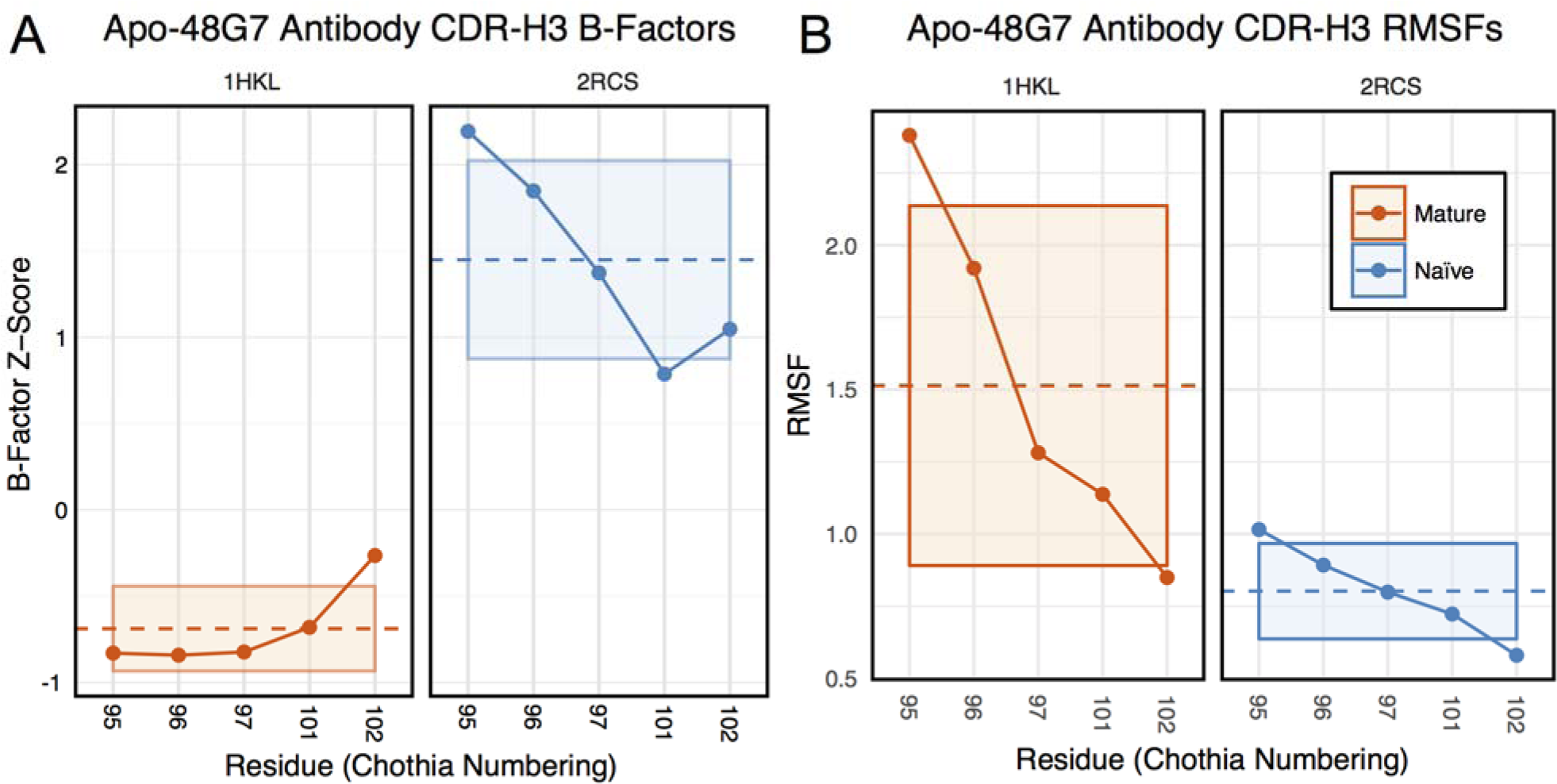
Analysis of catalytic antibody 48G7. (A) Comparison of normalized B-factor values for the CDR-H3 loop of the 48G7 antibody in crystal structures of the unbound naïve (dark blue) and mature (dark orange) antibodies. The dashed line indicates the average value and is outlined by a box defined by the average plus-or-minus the standard deviation. (B) Comparison of CDR-H3 loop RMSFs for the MD simulations of the naïve and mature 48G7 antibodies.

Prior follow-up studies on 48G7 used MD simulations to assess flexibility. Briefly, 500 ps short MD simulations of the naïve and mature antibodies in the presence of antigen with an explicit solvent model (TIP3P) found the CDR-H3 loop to be more flexible in the naïve than in the mature antibody by comparison of RMSFs (30), but 15 ns MD simulations of the naïve and mature antibodies in the absence of antigen with an implicit solvent model (GB/SA) found no difference between the two, again by comparison of RMSFs (32). Another study based on an elastic network model also suggested that, in the absence of antigen, the fluctuations of the naïve and mature 48G7 were similar, but their binding mechanisms could differ depending on response to antigen binding; the naïve antibody shows a discrete conformational change induced by antigen whereas the mature antibody shows lock-and-key binding where antigen reduce flexibility of the mature antibody (61). Due to the contentious nature of these results, we ran 200 ns MD simulations for the apo-48G7 naïve and mature antibodies in the absence of antigen with an explicit solvent model (TIP3P). We measured both RMSDs and RMSFs for the Cα atoms along the CDR-H3 loop and computed the difference between the naïve and mature antibodies (Supplementary Table 2). Figure 7B shows that the CDR-H3 loop RMSFs are consistently greater for the mature than the naïve 48G7 antibody.

Finally, as we have done through this study, we used FIRST-PG to measure CDR-H3 loop flexibility. To limit the effects of crystal structure artifacts on FIRST-PG analysis, we used an ensemble of ten representative structures, derived by clustering trajectory frames and selecting ten structurally distinct cluster medians from the MD simulations, similar to a previous flexibility study for this antibody (33). The CDR-H3 loop flexibility of apo-48G7, as determined by FIRST-PG analysis of MD ensembles is shown in Figure 8. The FIRST-PG analysis showed no significant difference between the mature and naïve antibodies.

**Figure 8.**
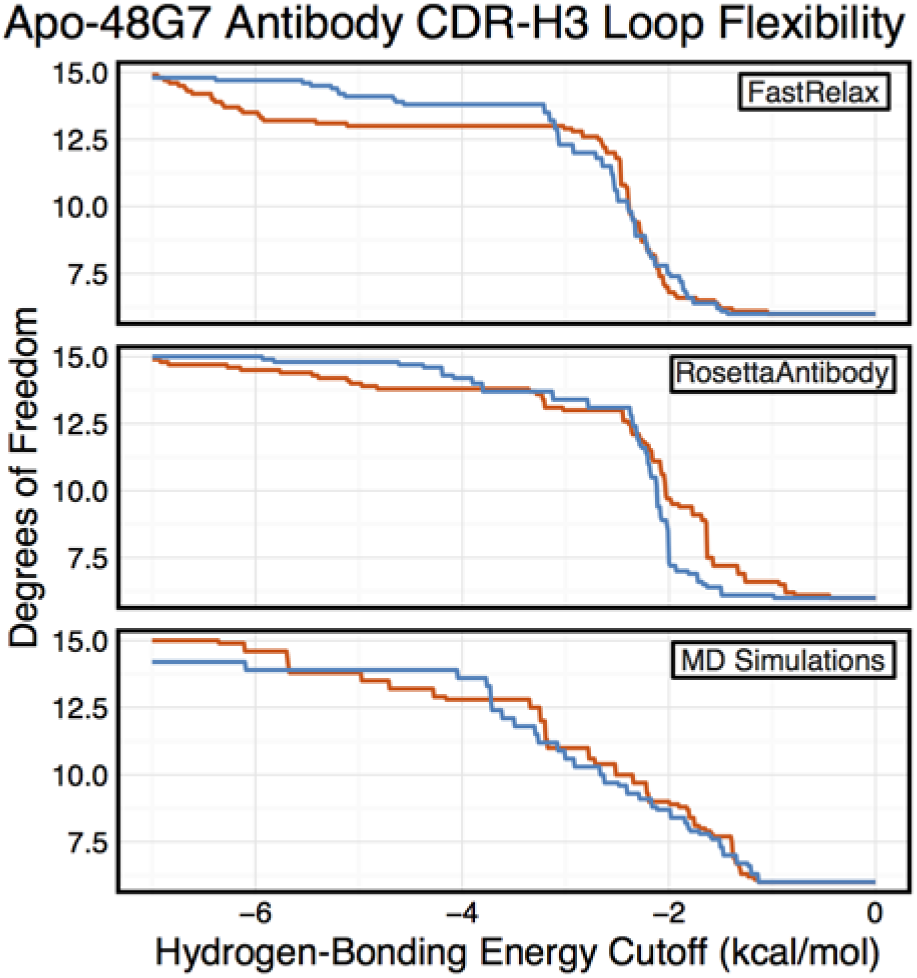
Comparison of FIRST-PG results for naïve (dark blue) and mature (dark orange) 48G7 antibodies using either Rosetta FastRelax, RosettaAntibody, or MD to generate 10-member ensembles. FIRST-PG analysis calculates the DOFs of CDR-H3 loop as a function of hydrogenbonding energy cutoff. FIRST-PG analysis of the FastRelax ensemble shows similar DOF counts in the range 0 to −3 kcal/mol for the naïve and mature antibodies, however, for higher energy cutoffs, the naïve antibody has more DOFs, at the same energy cutoff, than the mature antibody. The result is similar for the MD ensemble. On the other hand, FIRST-PG analysis of RosettaAntibody ensembles shows the mature antibody possessing slightly more DOFs than the naïve antibody at low-energy cutoffs, with the opposite being true at high-energy cutoffs.

In addition to using MD simulations to generate ensembles, we used ensembles generated by RosettaAntibody and Rosetta FastRelax, permitting direct comparison. The CDR-H3 loop flexibility of apo-48G7, determined by FIRST-PG analysis of FastRelax and Rosetta Antibody ensembles, is shown in Figure 8. The curves from FastRelax and the MD simulation are similar for low-energy cutoffs (e.g. in the range of 0.0 to −3.0 kcal/mol), with the naïve and mature DOFs being the same. These curves diverge at higher-energy cutoffs where the FastRelax curve shows a more flexible naïve antibody and the MD curve does not. The curve from RosettaAntibody ensembles differs from the two and shows a more flexible mature antibody at low-energy cutoffs and a more flexible naïve at high-energy cutoffs. For less visual and more quantitative comparisons, we computed the AUC of the DOF versus hydrogen-bonding energy cutoff plots (Supplementary Table 2). We find the AUC is only slightly greater for naïve than mature antibodies in the FastRelax and RosettaAntibody ensembles, with the naïve AUC reducing by only 3.9% and 0.2%, respectively, upon maturation. MD ensembles show the opposite outcome, with the mature antibody having 1.3% greater AUC than the naïve.

Further validation was carried out on two other previously studied antibodies and reported in the Supplementary Table 2 and Supplementary Figures 3 and 4. For the 4-4-20 antibody, antigen-bound structures were compared and the average mature B-factors were within a standard deviation of the naïve. For the influenza antibody, average B-factors were compared between an unbound naïve and a bound mature crystal structure, showing significant rigidification. However, results are conflated due to the lack of unbound crystal structures, as in bound structures antibody-antigen contacts artificially increases rigidity of the CDR-H3 loop. In contrast to B-factor analyses, FIRST-PG analyses yielded mixed results for these two antibodies. The 4-4-20 antibody was found to become more flexible upon maturation by FIRST-PG analysis of all but Rosetta KIC ensembles. The influenza antibody was found to become more rigid upon mature by FIRST-PG analysis of all but Rosetta FastRelax ensembles. Finally, we analyzed RMSDs and RMSFs from MD simulations and found that the mature 4-4-20 antibody has higher CDR-H3 loop RMSD, but lower RMSF, values than the naïve while the mature influenza antibody was found to have lower values for both (Supplementary Table 2). We consider the significance of these results and compare them in detail to past analyses of flexibility in the Discussion section.

## 4 Discussion

### 4.1 The Varying Effects of Affinity Maturation on CDR-H3 Flexibility

Affinity maturation, through a series of somatic hypermutation events and selection processes, can evolve a low-affinity, naïve antibody to bind an antigen with both high affinity and specificity (62). Elucidating the affinity maturation process is desirable to understand molecular evolution, develop antibody engineering methods, and guide vaccine development (63). Past studies have suggested that, with few exceptions (29, 64, 65), naïve antibodies are highly flexible and maturation leads to improved affinity and specificity through the optimization and rigidification of the antibody paratope, and in particular the CDR-H3 loop (22, 27, 28, 31–33). However, these studies have been limited, often focusing on a single antibody and assessing flexibility indirectly. We sought to test the generalizability of the rigidification-upon-maturation hypothesis. We were enabled by the large number of antibody structures in the PDB, homology models generated from high-throughput repertoire sequencing data, and the FIRST-PG method for rapid structural flexibility calculation to ask whether affinity maturation leads to CDR-H3 loop rigidification.

Unexpectedly, in a comparison of flexibility of repertoires, our data show little difference between naïve and mature antibodies: FIRST-PG calculations showed no difference for RosettaAntibody homology model ensembles of the most common naïve and mature antibodies in human peripheral blood cells. The same calculations showed no difference in CDR-H3 loop DOFs of crystal structures under two different refinement schemes (FastRelax and KIC). Even after accounting for the presence/absence of antigen, CDR-H3 loop B-factor distributions were the same for both mature and naïve antibody crystal structures. These results indicate that rigidification of the CDR-H3 loop does not always occur upon affinity maturation.

Since our observations did not indicate clear rigidification over two sets of antibodies, we considered the following possibilities: (1) comparison of different length CDR-H3 loops was unfair because longer loops are inherently more flexible, (2) comparison of different antibodies was unfair because different combinations of gene segments and V_H_-V_L_ pairs will result in different flexibilities, (3) mutations within CDR-H3 loop, which we could not identify for the PDB set because of the difficulty in D/J-gene alignments, may have modulated flexibilities of CDR-H3, (4) inaccuracies in the computational methods could preclude observation of rigidification, and (5) FIRST-PG-measured backbone DOFs are not a good measure of flexibility. To address the first concern, we analyzed loops of consistent length via B-factor and FIRST-PG (Figures 1B & 2B, Supplementary Figures 4 & 5). We found that, according to KS testing and when accounting for the presence/absence of antigen, B-factor distributions were not distinct for naïve and mature sets of antibodies with 10-residue CDR-H3 loops. We also found that FIRST-PG DOF AUCs of the naïve and mature sets of antibodies with the same length CDR-H3 loops were within a standard deviation for both RosettaAntibody, FastRelax, and KIC ensembles. So, even when accounting for length, mature antibodies are not significantly more rigid than naïve ones.

To address the concern that comparison of sets of antibodies originating from different V_H_ and V_L_ genes is unfair, we analyzed mature/naïve antibody pairs that had been previously studied and mature/naïve-reverted pairs that we generated with RosettaAntibody and analyzed by FIRST-PG (Figures 6–8, Supplementary Table 2). We found that CDR-H3 loop B-factors did not always indicate rigidification upon maturation and on one occasion we observed the reverse (Supplementary Figure 16). We also found that mature antibodies did not always become more flexible upon naïve reversion, but instead displayed a breadth of behaviors (Figure 6). So, when analyzing matched naïve/mature pairs, we do not see consistent rigidification upon maturation.

Our analysis of previously studied naïve/mature antibody pairs coupled with the earlier repertoire analysis should alleviate concerns that our flexibility results for the PDB set were strongly affected by our inability to align D/J-gene segments and thus consider mutations in the CDR-H3 loop. The previously studied pairs included CDR-H3 mutations and the repertoire set had antibody sequences determined by Illumina MiSeq sequencing with naïve/mature status assigned by the absence/presence of the CD27 cell-surface receptor. In both cases, the naïve and mature sequences were determined through the entire Fv, and flexibility analysis still revealed mixed results.

Finally, to address the concern that RosettaAntibody models may not be accurate enough to be useful for FIRST-PG calculations, we tested FIRST-PG on a range of structural ensembles with varying deviation from the crystal structure. We found no difference in the naïve vs. mature antibody CDR-H3 loop AUC of the FIRST-PG results, regardless of the ensemble generation method used (Figure 2 and Supplementary Figure 4). We also determined flexibility through alternative measure such as crystal structure B-factors and RMSFs in MD simulations. For both, affinity maturation was not found to have a consistent, rigidifying effect. Thus, even if model inaccuracies confound analysis, other data support the same hypothesis.

### 4.2 Comparison with Prior Results

Our analysis included several antibodies that have been the subject of previous flexibility studies, permitting a direct comparison (Supplementary Table 4 summarizes past studies). One of the most studied antibodies is the anti-fluorescein antibody, 4-4-20. Spectroscopic experiments measuring the response of a fluorescent probe (fluorescein) and MD simulations measuring Cα atom fluctuations suggested that somatic mutations restrict conformational fluctuations in the mature antibody (26, 28, 31). Our analysis of 4-4-20 was not as clear: we observed no significant difference in naïve vs. mature CDR-H3 loop crystallographic B-factors (Supplementary Figure 14) and found the mature antibody to be more rigid in FIRST-PG calculations only in the −2.0-0.0 kcal/mol range of hydrogen-bonding energy cutoffs (Supplemental Figure 15). Similar mixed results were observed by Li *et al.* (33) who used a Distance Constraint Model (DCM) to analyze flexibility in an ensemble of 4-4-20 conformations drawn from MD simulations. They found increases in structural rigidity of the CDR-H3 loop, as determined by the DCM, occurred upon affinity maturation, but these increases did not correspond to decreases in dynamic conformational fluctuations, as determined by RMSFs from MD simulations. Further studies artificially matured 4-4-20 by directed evolution, resulting in a femtomolar-affinity antibody, 4M5.3 (66), but the crystal structures of 4M5.3 and 4-4-20 were almost identical (the reported backbone RMSD is 0.60 Å) and thermodynamic measurements suggested that the affinity improvement was achieved primarily through the enthalpic interactions with subtle conformational changes (67). This observation was contradicted by Fukunishi *et al.* (68), who performed steered MD simulations to analyze the effects of the mutations on the flexibility of 44-20 and 4M5.3. By applying external pulling forces between the antibodies and the antigen along a reaction coordinate, they quantified the interactions and showed that, during the simulations, fluctuations of the antibody, especially the CDR-H3, and of the antigen were indeed larger in 4-4-20 than in the more matured antibody, 4M5.3 (68). Thus, there is some variation not only in our results, but also in the literature as to the effects of affinity maturation on 4-4-20.

Another set of well-studied antibodies are the four catalytic antibodies: 48G7, 7G12, 28B4, and AZ-28. In fact, the first crystallography studies to suggest rigidification of the CDR-H3 loop as a consequence of affinity maturation were performed on 48G7. Wedemayer *et al.* observed larger structural rearrangements upon antigen binding in the CDR-H3 loop for the naïve antibody than the mature antibody (Supplementary Figure 12 & 13) (16). Crystallization of the naïve unbound, naïve bound, mature unbound, and mature bound states for 7G12, 28B4, and AZ-28 revealed similar results (18, 19). Additionally, MD simulations of the four catalytic antibodies in implicit solvent were used to calculate CDR Cα atom B-factors (32). Wong ***et al.*** showed a decrease in mature CDR-H3 loop B-factors in three cases (7G12, 28B4, and AZ-28) whereas no significant difference was observed for 48G7 (see Figure 2 in Wong et al.). Furthermore, for 48G7, Li *et al.* used MD simulation to generate structural ensembles and DCM analysis to determine flexibility. They found that the mature CDR-H3 loop is more rigid than the naïve, according to DCM, but used an unusual loop definition that included five additional flanking residues (see Fig. 1 in Li *et al.*), making comparison challenging (longer loops will be inherently more flexible), and they observed increases in the mature CDR-H3 loop RMSFs (see Fig. 8 in Li *et al.)* (33). Our analysis of CDR-H3 loop B-factors showed rigidification for 48G7 and 7G12, but not for 28B4 and AZ-28 (Figure 7, Supplemental Figures 16 & 17). FIRST-PG analysis of FastRelax, RosettaAntibody, and MD ensemble for 48G7 showed slight to no rigidification (Figure 7). Finally, MD simulations for 48G7 showed no difference in naïve versus mature CDR-H3 loop flexibility as determined by FIRST-PG and revealed higher RMSFs for the mature loop. Our mixed results for the effects of affinity maturation on 48G7 are consistent with literature, but there is variation between our results and the literature as to the effects of affinity maturation on the other catalytic antibodies.

Finally, Schmidt *et al.* used X-ray crystallography, MD simulations, and thermodynamics measurements to investigate how somatic mutations affected the binding mechanism of antiinfluenza antibodies (22). They identified three mature antibodies, their unmutated common ancestor (UCA), and a common intermediate, all derived from a subject immunized with an influenza vaccine. The affinities of the mature antibodies were about 200-fold better than the UCA. MD simulations of the UCA and the mature antibodies showed that CDR-H3 loop of the UCA could sample more diverse conformations than the mature antibodies, whose CDR-H3 loop sampled only conformations optimal for antigen binding, supporting the hypothesis that somatic mutations rigidify antibody structures. In another study by the same group (69), further MD simulations were performed on the same systems, showing that, although many somatic mutations typically accumulate in broadly neutralizing antibodies during maturation, only a handful of mutations substantially stabilize CDR-H3 and hence enhance the affinity of the antibodies for antigen. In our studies, all the results for the anti-influenza antibody, except FIRST-PG flexibility calculations for the Rosetta FastRelax ensemble, show rigidification of the CDR-H3 loop as an effect of affinity maturation and are in agreement with the detailed analysis of Schmidt *et al*.

For these three antibody families we analyzed in detail, we observed mixed effects of affinity maturation on two (catalytic antibodies and 4-4-20) and clear rigidification in one (anti-influenza antibody). For the two with mixed results, we note that past work has also shown conflicting results. We interpret these results as supportive of our repertoire-wide analysis that affinity maturation does not always rigidify the CDR-H3 loop.

### 4.3 Biophysical properties underlying antibody binding

Why is antibody CDR-H3 loop rigidification not a consistent result of affinity maturation? Consider the process of affinity maturation, which selects for antibody-antigen binding and against interactions with self or damaged antibodies (i.e. when deleterious mutations are introduced by activation-induced cytidine deaminase) (70). Under these selection pressures, what is the benefit of CDR-H3 loop rigidification? Loop rigidification can only decrease the protein-entropy cost for antibody-antigen binding, having ostensibly no effect on enthalpy and solvent entropy of binding, and self-interactions. If CDR-H3 loop rigidification is just one of many biophysical mechanisms that can be selected for during affinity maturation, then we do not expect to observe it consistently, in line with our results.

What are the other possible mechanisms then? Surprisingly, mutations leading to multi-specificity or promiscuity may be beneficial to selection: antibodies are multivalent, so an antibody capable of binding multiple antigens with intermediate affinity can gain an effective advantage through cooperative binding over an antibody capable of binding only one antigen. Unsurprisingly, multispecific mature antibodies have been observed. Take for example the anti-hapten antibody, SPE7 (71). Crystal structures of SPE7 with different antigens and in its apo-state demonstrate that SPE7 can assume different conformations. Motivated by these observations, Wang *et al*. exploited MD simulations to investigate the binding mechanisms of SPE7 (72). The MD simulations and subsequent analyses suggested that multi-specific antigen binding is mediated by a combined mechanism of conformer selection and induced fit. This behavior could not have arisen if CDR H3 loop rigidification were a consistent result of affinity maturation.

## 5 Conclusions

We have conducted the largest-scale flexibility study of antibody CDR-H3 loops, analyzing ~1,000 crystal structures and ~2,000 homology models. We used B-factors and FIRST-PG to assess flexibility. We sought to identify the effects of affinity maturation on CDR-H3 loop flexibility, expecting the CDR-H3 loop to rigidify. We found that there were no differences in the CDR-H3 loop B-factor distributions or FIRST-PG DOFs for naïve vs. mature antibody crystal structures and in the CDR-H3 FIRST-PG DOFs for homology models of repertoires of naïve and mature antibodies. These findings suggest that there is no general difference between naïve and mature antibody CDR-H3 loop flexibility in repertoires of naïve and mature antibodies. However, we observed rigidification of the CDR-H3 loop for some antibodies when the mature antibody was compared directly to its germline predecessor. So, it is possible that increased rigidity occurs alongside other affinity-increasing changes. We conclude that pre-configuration of the paratope (which typically contains the CDR-H3 loop) is just one of many mechanisms for increasing affinity.

Further work must be done to address the issues observed here, *i.e.* inconsistent results across the different methods used to measure flexibility. One possible route is to explore experimental methods that directly measure protein dynamics across several timescales, and use them to study a relatively large (more than one or two antibodies) and diverse (e.g. from different source organisms or capable of binding different antigens) set of antibodies. For example, HDX-MS is capable of identifying protein regions with dynamics on timescales from milliseconds to days (73).

Finally, we note the need for more rapid and accurate antibody modeling methods. With the advent of high-throughput sequencing, there now exits a plethora of antibody sequence data, but little structural data. Accurate modeling can overcome the lack of high-throughput structure determination method and provide crucial structural data. These structures can then be used to address scientific questions on a larger scale than before, on the scale of the human antibody repertoire.

## 7 Author Contributions

JJ, AS, and JG designed the research. JJ, AS, DK, and NT performed the research. JJ, AS, and DK analyzed the data. JJ, AS, DK, NT, NK, KT, and JG wrote the paper.

## 9 Acknowledgments

The authors would like to acknowledge Oana I. Lungu and Erik L. Johnson for sharing the antibody repertoire homology models. The super-computing resources in this study have been provided in part by the Maryland Advanced Research Computing Center, the ROIS National Institute of Genetics, and the Human Genome Center at the Institute of Medical Science, The University of Tokyo, Japan.

